# Machine Learning Meets Pharmacokinetics: A Comparative Analysis of Predictive Models for Plasma Concentration-Time Profiles

**DOI:** 10.1101/2025.05.05.652167

**Authors:** Felix Jost, Clemens Giegerich, Christoph Grebner, Hans Matter, Henrik Cordes

**Affiliations:** Sanofi R&D, Translational Medicine Unit, Quantitative Pharmacology, Research Pharmacometrics, Frankfurt, Germany; Sanofi R&D, Translational Medicine Unit, Quantitative Pharmacology, Disease Modeling, Frankfurt, Germany; Sanofi R&D, Integrated Drug Discovery, Frankfurt, Germany

## Abstract

Predicting pharmacokinetic (PK) profiles from molecular structures constitutes a significant advancement in pharmaceutical research with substantial implications for expediting the entire drug discovery process. Our investigation presents a comprehensive comparative analysis of five distinct methodological frameworks for predicting rat plasma concentration-time profiles: four approaches that integrate mechanistic modeling with computational techniques, and one approach employing pure machine learning (ML) architecture without physiological constraints. As reference (1), we used a method that predicts non-compartmental analysis (NCA) parameters to predict plasma PK profiles with a one-compartmental PK model (NCA-ML). Alternative approaches include: (2) a physiologically based PK (PBPK) model utilizing ML-predicted *in vitro* characteristics; (3) CMT-ML, where neural networks predict compartmental PK model parameters; (4) CMT-PINN, employing physics-informed neural networks; and (5) PURE-ML, using decision trees to predict concentration values at specific timepoints based on predicted volume of distribution and clearance. Assessment using multiple performance metrics demonstrated that the CMT-PINN approach achieved superior predictivity for PK profiles. Models trained directly on concentration-time, rather than derived PK parameters, delivered markedly improved predictive performance, particularly when working with limited training datasets. Our findings confirm the viability of predicting PK behavior from molecular structures before synthesis, representing a significant advancement for subsequent design. Implementation of these computational approaches enables informed compound selection early in the discovery pipeline, concentrating valuable resources for advanced preclinical investigations on only the most promising candidates, with anticipated beneficial effects on overall development timelines.

## INTRODUCTION

Pharmaceutical research aims to enhance patient lives through innovative treatments. The initial drug discovery phase focuses on identifying promising molecules for development into effective, marketable medications. The primary challenge lies in efficiently narrowing down drug candidates with an optimal balance of *in vitro* and *in vivo* properties. To address this, modern drug discovery employs a sophisticated pipeline that strategically eliminates unsuitable compounds early through a staged approach. This process utilizes an increasingly complex profiling tree, enabling data-driven decisions at each progression step.

Drug discovery projects typically involve a combination of high throughput screening and computational approaches to identify interesting molecules with specific physicochemical properties, target binding, and ADME characteristics (absorption, distribution, metabolism, elimination). Promising compounds then undergo comprehensive profiling in specialized assay panels that evaluate their potency, physicochemical properties, and ADME profile in greater detail. Selected molecules are subsequently synthesized at larger scale for evaluation in sophisticated *in vivo* studies examining pharmacokinetic (PK), pharmacodynamic (PD), and preliminary safety (PS) parameters in animal models. This staged approach optimizes resource allocation in candidate selection. Increasingly, data-driven technologies are revolutionizing this workflow, as validated machine learning (ML) models can potentially replace certain costly and time-intensive experimental procedures with rapid computational predictions.

In the assessment of PK profiles, the labor-intensive and expensive *in vivo* experiments represent the primary workflow constraint within the discovery cascade. Those studies provide drug concentration-time profiles in blood plasma, target organs, and other tissues – and therefore represent the aggregated result of all bio-chemical and –physical interaction processes of a molecule entering a body. These exposure profiles can be represented by various mathematical models, such as non-compartmental analysis (NCA), compartmental PK modeling (CMT), physiologically based PK models (PBPK) and others.^1^ Parameters of such models in combination with scaling or translational approaches such as *in vitro / in vivo* correlation (IVIVC), allometric scaling or PBPK modeling are utilized to estimate the PK profile in humans. From an ensemble of suitable molecules after testing, ideally representing different chemical series, the most promising one is selected for clinical development showing an optimal balance of its PK, PD, and PS profiles.

Within healthcare and drug discovery,^2^ particularly during early target identification and hit finding phases,^3^ cutting-edge computational approaches for analyzing complex, heterogeneous datasets offer opportunities to enhance workflow efficiency. Research organizations are increasingly embracing a paradigm where a molecule’s complete profile can be computationally predicted, shifting emphasis from extensive data generation toward a prediction-validation framework. This transformation is exemplified by the widespread adoption of “design-make-test-analyze” (DMTA) cycles that systematically leverage accumulated knowledge and experimental data, reflecting a fundamental transformation in discovery methodology.

To this end, novel technical advancements aim to accelerate multidimensional optimization of chemical series while reducing the amount of animal studies. An ever-increasing array of ML approaches has been developed and is nowadays used to predict *in vitro* ADME, target properties and *in vivo* PK characteristics based on the numerical representation of molecular structures.^4^ Ideally, such methods are integrated in compound design prior to time-consuming synthesis and testing. Many contributions have been made over the past years to facilitate these developments regarding method development,^5–7^ and implementation.^8^ Due to the multitude of properties and processes resulting in PK profiles, any prediction is still challenging, and recent approaches focus on a defined parameter space limited by the species, dose range, administration routes, data set sizes and chemical space to evaluate their methods.

Most studies employ model-based (hybrid) PK prediction approaches that vary in the employed ML and mechanistic modeling approach. Such approaches are often trained using NCA parameters derived from existing PK profiles as dependent variables, such as clearance (CL), half-life (t1/2), and volume of distribution at steady state (Vss). When combining these NCA parameters with one-compartment models for analysis (NCA-ML), a monophasic PK time-concentration curve can be predicted.^9^ Other approaches (CMT-ML) employ multi-compartment or whole-body PBPK (PBPK-ML) models, nowadays also parameterized with ML predicted physicochemical and *in vitro* assay properties.^9–16^ Approaches also directly use PK profiles as input for training and prediction of PK profiles via ML techniques with (CMT-PINN) and without (PURE-ML) utilizing mechanistic models.^17–20^ Here, challenges include the interpretability (PURE-ML) and the prediction quality of complex methods (e.g. PBPK-ML) due to the combination of *in vitro* and *in vivo* data. The combination of multiple data types and sources with varying accuracy includes an additional layer of compacity with respect to predictivity and translatability.^21^ Nevertheless, recent contributions have shown that predicted *in vitro* characteristics translate to *in vivo* PK descriptions.^10^ A practical compromise involves defined compartmental models (one-, two-, and three-compartments) that provide adequate data description comparable to mechanistic models with less mathematical equations and model assumptions. Here, the underlying aggregated ADME processes are imprinted as bias in the model structure resulting in the to-be-expected characteristics of empirically observed exponential concentration declines in plasma PK profiles. The observed exponential decay is routed in the first order rate of the underlying biochemical reaction, where under substrate/reactant limiting conditions, the reaction rate is proportional to the concentration.

Recent review articles have summarized various prediction methods for PK across different animal species,^22^ and human data.^23^ These methods rely on plasma PK profiles for which standardized public repositories are sparse.^24,25^ However, these studies differ significantly with respect to methodological details, data sets, species investigated, generation of test and validation sets, and performance metrics. This makes a comparison across different approaches challenging. Table 1 provides an overview of methods and characteristics recently published in the literature (until the end of 2024).

**Table 1:**
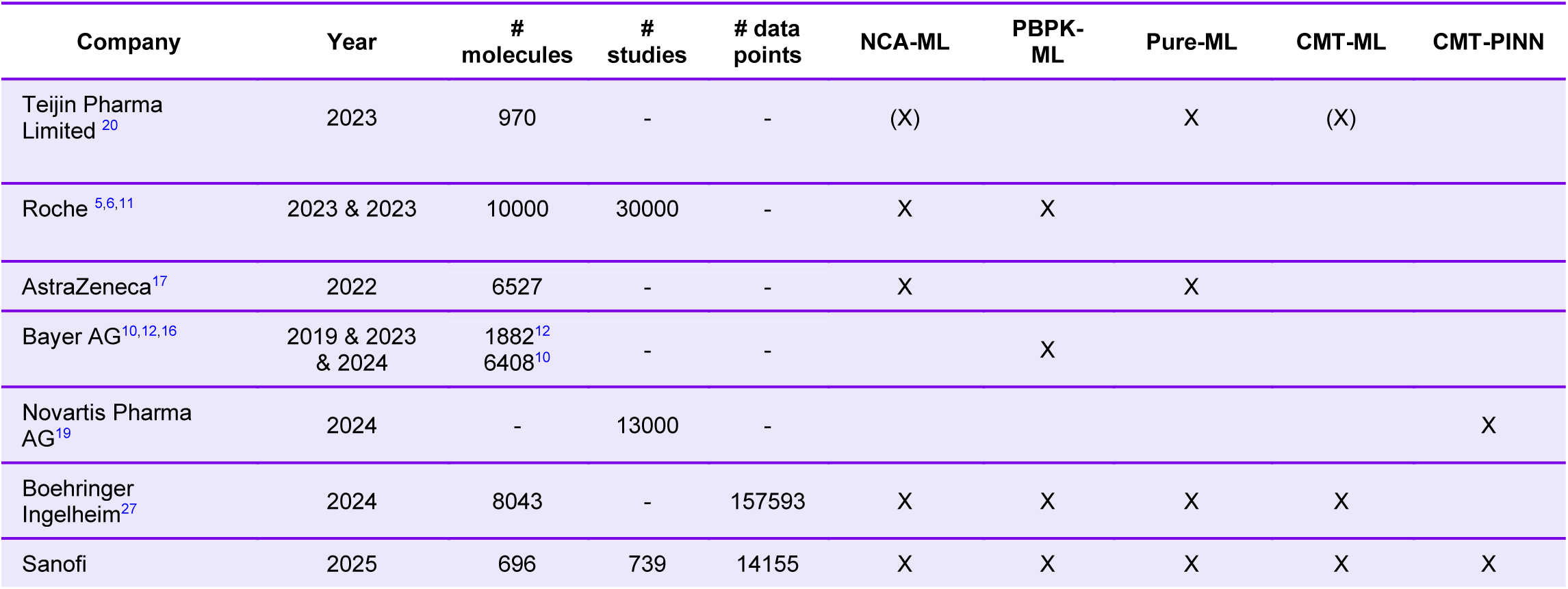
Overview on recently published AI/ML approaches to predict PK profiles. “X” indicates, which prediction method was investigated and “-“specifies, if specific data information was not disclosed.

| Company | Year | # molecules | # studies | # data points | NCA-ML | PBPK-ML | Pure-ML | CMT-ML | CMT-PINN |
| --- | --- | --- | --- | --- | --- | --- | --- | --- | --- |
| Teijin Pharma Limited<sup>20</sup> | 2023 | 970 | - | - | (X) |  | X | (X) |  |
| Roche<sup>5,6,11</sup> | 2023 & 2023 | 10000 | 30000 | - | X | X |  |  |  |
| AstraZeneca<sup>17</sup> | 2022 | 6527 | - | - | X |  | X |  |  |
| Bayer AG<sup>10,12,16</sup> | 2019 & 2023 & 2024 | 1882<sup>12</sup><br>6408<sup>10</sup> | - | - |  | X |  |  |  |
| Novartis Pharma AG<sup>19</sup> | 2024 | - | 13000 | - |  |  |  |  | X |
| Boehringer Ingelheim<sup>27</sup> | 2024 | 8043 | - | 157593 | X | X | X | X |  |
| Sanofi | 2025 | 696 | 739 | 14155 | X | X | X | X | X |

Towards an objective comparison across multiple approaches, we compared four model-based hybrid methods and one ML method to predict *in vivo* rat plasma PK profiles directly from molecular structures employing a consistent data set. As reference, we use a hybrid prediction method based on NCA parameters together with a one-compartmental PK model for predicting concentration-time profiles (NCA-ML). All other approaches are compared to this method. The second hybrid method uses ML predicted *in vitro* characteristics which serve as input for a PBPK model to predict the *in vivo* PK profiles (PBPK-ML). The two other hybrid methods range between approaches using NCA and mechanistic PBPK models. In both approaches a neural network (NN) is trained to predict parameters of a compartmental PK model, which then generates concentration-time profiles. The approaches differ in the underlying training methodologies (CMT-ML & CMT-PINN). In CMT-ML a neuronal network (NN) is trained on estimated PK parameters from one-, two-, and three-compartmental models, whereas in CMT-PINN the NN is directly trained on the concentration-time profiles. This latter approach is known as a physics-informed neural network (PINN).^26^ Both methods allow for interpretation of PK parameters, demonstrate good predictivity of PK profiles, and were investigated in different reports.^19,27^ Besides hybrid models, the last approach is a pure ML approach (PURE-ML) which is also trained on PK profiles, but without using a compartmental PK model.

Our objective is to establish a standardized evaluation framework across diverse ML approaches using a consistent dataset, consistent data partitioning, and uniform performance metrics, thereby enabling direct methodological comparisons. This also allows to put our results in perspective to recent literature on prediction of PK concentration-time profiles from molecular structures alone.

## METHODS

### Characteristics of used data set & preparation

We selected experimental data from a total of 739 study arms across 721 *in vivo* rat PK studies after intravenous administration for 696 different molecules at Sanofi. The data set was curated such that only PK profiles under non-limiting conditions were used excluding analytical outliers, animals showing incorrect administration and high doses and resulting in a total of 14,155 plasma PK data points. The administered dosages range from 0.1 to 10 mg/kg with 89% of the studies using 1 mg/kg and 9% using 3 mg/kg. Typically, each study arm consisted of 2 (83%) to 3 (15%) animals, with sequential sampling conducted up to 24 hours post-dose. Notably, the plasma PK sampling is denser during the first hour (2 to 5 samples), then switched to every second hour from 2 to 8 hours (1 to 6 samples) and concluded at 24 hours post-dose (0 to 1 samples). All ML approaches use the same underlying PK data set.

### Used splitting strategy

The prepared data set was divided into training, validation, and test sets with a split of 80%, 10%, 10% using a chemical diversity-based algorithm (Min-Max-Picking, as implemented in RDKit^28^). All ML approaches use the same data split.

### Metrices

We assessed model prediction accuracy with a comprehensive set of statistical metrics across multiple dimensions, incorporating measures of central tendency (mean, median) and dispersion (variance) including coefficient of determination (R^2^), mean absolute percentage error (MAPE) and median absolute percentage error (MAPE) on log-transformed data (-log), Spearman’s correlation coefficient, median fold change error (MFCE) and geometric mean fold error (GMFE) between estimated and predicted concentration vs. time data. This latter evaluation metric does not rely on observed data, which would bias any method comparison towards early PK sampling time points due to a more frequent sampling compared to the late time points. In contrast, this metric considers all phases of PK time series equally. Via the usage of estimated data also a comparison between publications can be performed. Estimated concentration data is generated with the corresponding compartmental model utilizing the estimated PK parameters (depending on number of compartments: V1, V2, V3, CL1, CL2, CL3). The selection of PK model for each compound to generate the estimated concentration data is described in detail in the next section *CMT-ML* and in literature.^27^

### Overview approaches

In this work we developed, evaluated, and compared five different model-based ML approaches to predict plasma concentration-time profiles directly from molecular structures. Here, three prediction approaches naming PURE-ML, NCA-ML, and CMT-ML need a preparation step. The PURE-ML and NCA-ML approaches require an initial NCA, while the CMT-ML approach dedicated one-, two-, and three-compartment PK parameter estimations. For the NCA based methods, the analysis was done for a mean concentration-time profile for each study and subsequent dose group. For the CMT-ML approach, the PK model result in the lowest dose group or with the lowest Akaike Information Criterion (AIC) value was selected. The mean concentration-time profiles were also used for the training of the PURE-ML approach. The fourth approach, CMT-PINN, does not require additional preparation steps and training can be performed directly on the concentration-time profiles. The fifth approach, PBPK-ML, uses predicted *in vitro* characteristics which serve as input for a PBPK model to predict the *in vivo* PK profiles. A visual representation of the five approaches is shown in Figure 1.

**Figure 1:**
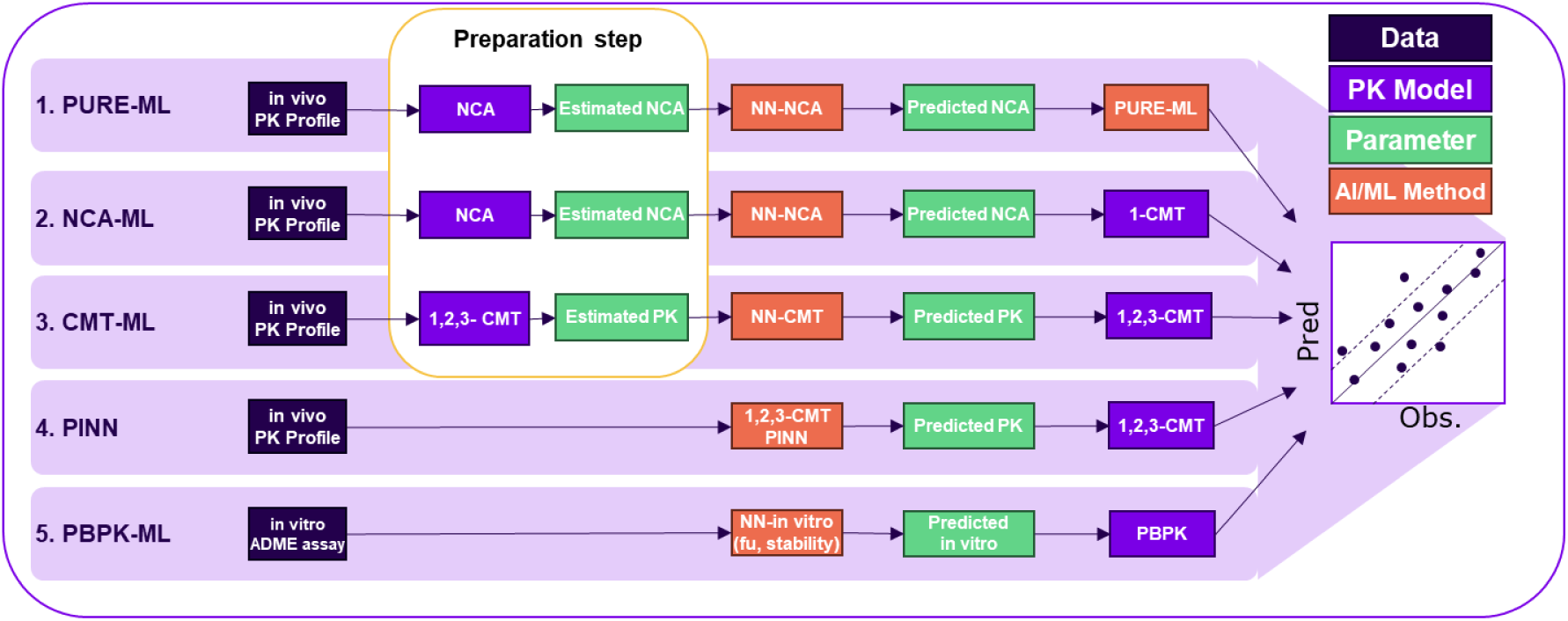
Schematic representation of the five tested approaches for predicting plasma pharmacokinetic profiles.

### Description of all approaches in detail

#### PURE-ML

The PURE-ML approach requires an initial preparation step in which NCA parameters (Vss and CL) are estimated from PK profiles. Next, a prediction method is developed that predicts NCA parameters from molecular structures (see next section). Finally, the method trains on the logarithmized concentration-time data while using the predicted NCA parameters and the administered dose as input values (see ^18^ for details). Instead of an explicit PK model, the trained decision trees predict concentration values at certain time points. The predicted concentration time points can then be used to derive a PK profile.

#### NCA-ML

The NCA-ML approach is based on the PK parameters Vss and CL. In a preparation step, the experimental PK profiles are used to calculate the NCA parameters Vss and CL. The NCA parameter calculation was performed for each compound across individual dose groups of every study resulting in a single Vss and CL value for each compound. Next, a graph convolutional neural network (GCNN) is trained on the calculated NCA parameters with the molecular structures encoded as Simplified Molecular Input Line Entry System (SMILES) string and transformed to a numerical representation as only input.^29^ The NCA-ML predicted NCA parameters are used as input for an one-compartmental PK model to simulate a monophasic PK profile to evaluate the overall prediction performance with respect to the experimental PK profiles. This NCA-ML approach serves as reference for comparison.

#### CMT-ML

The CMT-ML approach works like the NCA-ML approach. A preparation step is required in which experimental PK profiles are used to sequentially estimate one-, two-, and three-compartment PK model parameters (V1, V2, V3, CL1, CL2, CL3). This approach enables multiphasic PK profile description. For each compound, study, and dose group a one-, two-, and three-compartment model is fitted to the respective PK profiles. For each PK model parameter estimation previous NCA results are used as initials. Notably the individual PK profiles and a population PK approach with interindividual variability on all parameters and an exponential error model are used for parameter estimation.^30^ The lowest AIC is used as decision criteria for selecting the final PK model and respective parameters for further processing. In addition, a coefficient of variation < 70% for all model parameters is used as quality criteria for identifiability of parameters. Next, a GCNN is trained on the estimated PK parameters like in the NCA-ML approach in which each compound has a set of parameters V1, V2, V3, CL1, CL2 and CL3. Mapping of one– and two-compartmental models to a three-compartment model is enabled by setting the missing parameters V2 and V3 to a small value (0.1 L/kg) and CL2 and CL3 to a high value (12 L/ (h kg)) such that the three-compartment model behaves as a one or two compartment model. Finally, the trained GCNN provides predicted PK parameters as input for a three-compartmental PK model to simulate a multiphasic PK profile that is used to evaluate the overall prediction performance compared to experimental PK profiles (3CMT-ML). Additionally, we evaluated this approach for a setting with one– and two-compartmental models (1CMT-ML and 2CMT-ML) only to reduce the multitask ML architecture from six to four PK parameters as prediction method output.

#### CMT-PINN

The third approach CMT-PINN uses the same underlying PK models and evaluation method as the CMT-ML approach. However, instead of predetermining the best PK parameters, their estimation is part of the PINN model training itself and thus carried out directly on the concentration-time profiles (compare Figure 1). Here we evaluated two CMT-PINN settings, one that builds on one– and two-compartmental models only (2CMT-PINN) and one setting in addition includes a three-compartmental model (3CMT-PINN).

#### PBPK-ML

The PBPK-ML approach builds upon ML models for the prediction of physicochemical compound properties (logD, molecular weight, pka, pkb, halogen content) and early *in vitro* ADME assay parameters (microsomal stability, plasma protein binding). The set of parameters is predicted for each compound and used to build a whole body PBPK models for each study and dose group using a mean rat individual with the default weight generated from the PK-Sim physiology database and used for all simulations. Clearance of the compound was defined as total hepatic clearance “liver plasma clearance” in PK-Sim. Renal elimination was defined as glomerular filtration with “GFR fraction” set to 1 as default, no further active processes contributing to distribution or metabolization were considered.

Next, ten individual models accounting for different combinations of calculating cellular permeabilities and partition coefficients (two permeability, five partition model) are used to predict plasma PK profiles. The model performance was assessed by comparing predicted with experimental PK profiles.

### Machine learning architectures

In this section for each approach the NN architecture and the training procedure is described in detail.

#### PURE-ML

The PURE-ML approach employs extreme gradient boosting (XGBoost) as it was identified as best performing ML method compared to linear regression, Poisson regression, support vector machine and light gradient boost machine.^18^ For hyperparameter optimization of the XGBoost method 150 parameter combinations are used from the following sets: Max depth: {2,3,6,10,15}, learning rate: {0.001,0.005,0.01,0.1,0.3,0.5} and number of estimators: {10,25,50,100,200}. As input, the predicted Vss and CL values from NCA-ML are used together with the administered dosage. The same data set split was used among the different approaches to enable comparability. During the grid search, the configuration with the smallest validation loss was chosen as the best performing model and later evaluated on the test set.

#### NCA-ML & CMT-ML: Graph convolutional neural networks (GCN) for PK parameters from NCA and compartmental modeling

For the NCA-ML and CMT-ML, three multitask GCNNs were developed to predict the NCA parameters (Vss, CL), PK model parameters (V1, V2, V3, CL, CL2, CL3), and PK model parameters (V1, V2, CL, CL2), respectively.^31^ For CMT-ML, only the population parameter values were used to train the NN model.

All parameter values were logarithmically transformed (log_10_) before training. Individual models were trained on minimizing L2-loss (sum of squared differences between observed and predicted values) and by monitoring the Pearson correlation score (PCS) for the training data set and evaluated on the validation data set. For hyperparameter tuning, various models with different hyperparameter combinations were built. The best performing parameter combination regarding PCS on the validation set was used for further calculations. Training was stopped, if the loss of PCS did not improve for 20 subsequent steps (early stopping).^3^ For all GCNN calculations a previously reported implementation was used.^7^

#### CMT-PINN: Physical informed neural networks

Three PINNs were set up either using a one-, two-, or three-compartmental PK model. As loss function, the equally weighted logarithmic difference between concentration-time observations and predictions were divided by the number of samples and the number of individuals. As for the earlier approaches the same data set split was used for the PINNs to enable later comparability. For each of the three settings, the number of hidden layers (3, 4, 5, 6), the number of nodes (32, 64, 128, 256, 512, 1024), alternative activation functions (r tanh, sigmoid, relu, leakyrelu, elu, celu, rbf), the batch sizes (32, 64, 128) for the minibatch training, and different learning rates (0.0005-0.01) with Adam and AdamW were evaluated.

#### PBPK-ML: Graph convolutional neural network for *in vitro* parameters

For the PBPK-ML approach compound properties (molecular weight, halogen content, pka, pkb) and internally available prediction methods (lipophilicity, plasma protein binding, microsomal stability) were used to build bottom-up whole-body PBPK models without further optimization on parameters, scaling or distribution models. The used Sanofi internal global *in silico* ML models are generated and routinely applied in different discovery scenarios to profile virtual chemical structures after design to support decision making on the synthesis. These global models are derived from novel, consistent and harmonized experimental data and were described earlier in the literature.^7^

### Software

NCA, compartmental modeling, PBPK modeling, descriptive statistics and visualization was performed within RStudio^32^ (version 4.2.0). NCA was performed with the package PKNCA^33^ (version 0.10.2). For parameter estimation, the Stochastic Approximation Expectation Maximization (SAEM) algorithm implemented in nlmixr2^34^ (version 2.0.8) was used. PBPK modeling was performed with the ospsuite-R package^35^ (version 12.1.0) providing functionalities of loading, manipulating, and simulating *.pkml* model files. The simulations were created in the Open Systems Pharmacology Suite, Version 11.1 (REF: PMID 31671256) and exported to *.pkml* model files.

The ML calculations for the GCNN were performed using an inhouse python toolkit based on DeepChem for ML as described elsewhere^3^ and RDKit^28^ (version 2021.03.3) for molecular structures using internal scripts for data preparation, data set splitting, training, and prediction.

The PURE-ML method in form of XGBoost was used in Python as well with the package xgboost^36^ (version 1.7.6.).

The PINNs were set up in Julia^37^ (v1.10.4) with Flux^38^ (v0.14.19) using the optimization solvers Adam and AdamW and the native Vern7() ODE solver from OrdinaryDiffEq^39^ (v6.89.0). A sparse version of RDKit^40^ (v1.2.0) is available in Julia for calculating Morgan fingerprints.^29,41^

## RESULTS

Firstly, the best ML architectures after hyperparameter optimization are presented. For the PURE-ML approach, the best performing hyperparameters were identified as a maximal depth of 6, a learning rate of 0.1, and total number of estimators as 100. The final model for NCA-ML consists of two graph layers with 128 nodes, one final dense layer with 256 nodes, a learning rate of 0.0005 and no drop out mask. The CMT-ML model consists of three graph layers with 128 nodes, one final dense layer with 128, a learning rate of 0.001 and a drop out probability of 10%.

The best performing CMT-PINN was achieved with three full connected hidden layers (with size of 128 for one– and two-compartment models, for the three-compartment model the first hidden layer was increased to 1024), dropout layers (dropout rate of 0.08 for the one-, 0.085 for the two– and 0.125 for the three-compartment model), relu activation function, softplus output activation function, batch size 32, epochs 1000 and the default learning rate 0.001 of Adam optimizer.

Next, we evaluated the performance of multiple ML approaches (PURE-ML, NCA-ML, CMT-PINN, PBPK-ML, and CMT-ML) to predict plasma concentration-time data in rats using a test dataset comprising 70 compounds, except for CMT-ML which included 63 compounds due to parameter identifiability filtering criteria based on the coefficient of variation (see method section and Supplement table 1). We used a set of different statistical performance metrics (mean, median, variance, R2, MAPE, MAPE, MFCE, GMFE), common PK parameters (AUC, C_0_, C_min_), as well as the individual concentration-time profiles to benchmark the tested prediction algorithms.

Among all tested methods, the CMT-PINN approach performed best across all tested metrics (Table 2). It achieved the highest explainable variance with a R²-log of 0.854 and Spearman correlation of 0.933, along with the lowest error metrics (MAPE-log: 24.5%, MDAPE-log: 8.64, MFCE: 1.02). The PURE-ML approach followed closely with lower performance metrics (R²-log: 0.789, Spearman: 0.896) but achieved the lowest MAPE-log value at 22.1%. In contrast, NCA-ML and CMT-ML showed substantially lower prediction performances (NCA-ML: R²-log: 0.444, Spearman: 0.865; 3CMT-ML: R²-log: 0.471, Spearman: 0.852) with considerably higher error metrics. The NN for NCA-ML (mean-Pearson r² score: 0.68) outperformed the CMT-ML approach (2CMT-ML: 0.54, 3CMT-ML: 0.38) in prediction accuracy. PBPK-ML showed variable performance with R²-log ranging from 0.284 to 0.639 and Spearman values between 0.532 and 0.842, depending on the used permeability and partitioning coefficient estimation model.

**Table 2:** Summary of evaluation metrices on the test data set for NCA-ML, 3CMT-ML and 3CMT-PINN. CMT-ML and CMT-PINN use a three compartmental model.

| Approach | R2-log | Spearman | MAPE-log [%] | MDAPE-log | MFCE |
| --- | --- | --- | --- | --- | --- |
| **NCA-ML** | 0.444 | 0.865 | 96.6 | 13.6 | 1.26 |
| **PBPK-ML** | 0.284-0.639 | 0.532-0.842 | 50.5-78.6 | 20.8-33.9 | 0.223-0.596 |
| **3CMT-ML** | 0.471 | 0.852 | 90.3 | 15.9 | 0.870 |
| **PURE-ML** | 0.789 | 0.896 | 22.1 | 9.83 | 1.12 |
| **3CMT-PINN** | 0.854 | 0.933 | 24.5 | 8.64 | 1.02 |

R2-log = Pearson $r^2$ on log-transformed data, MAPE-log: Mean absolute percentage error on log-transformed data, MDAPE-log: Median Absolute
Percentage Error on log-transformed data, MFCE: mean fold change error

When examining the percentage of predicted concentration data points within acceptable error margins (Table 3), CMT-PINN again achieved the best performing results with 65.9% of predictions within two-fold error and 83.5% within three-fold error of observed concentrations. PURE-ML followed closely with 61.0% within two-fold and 79.7% within three-fold error. The other hybrid methods showed substantially lower inclusion rates: NCA-ML (11.6%/18.0%), 2CMT-ML (7.17%/11.5%), 3CMT-ML (8.98%/14.3%), and PBPK-ML (18.6-27.8%/30.9-45.1%). This performance pattern is reflected in the goodness-of-fit plots between observed and predicted concentrations (Figure 2). NCA-ML and CMT-ML notably show left-skewed correlation, constantly underpredicting lower concentration values. These lower values are typically seen from later measurement time points of the PK profile, indicating the limitation of these approaches to appropriately capture the PK profile shape.

**Figure 2:**
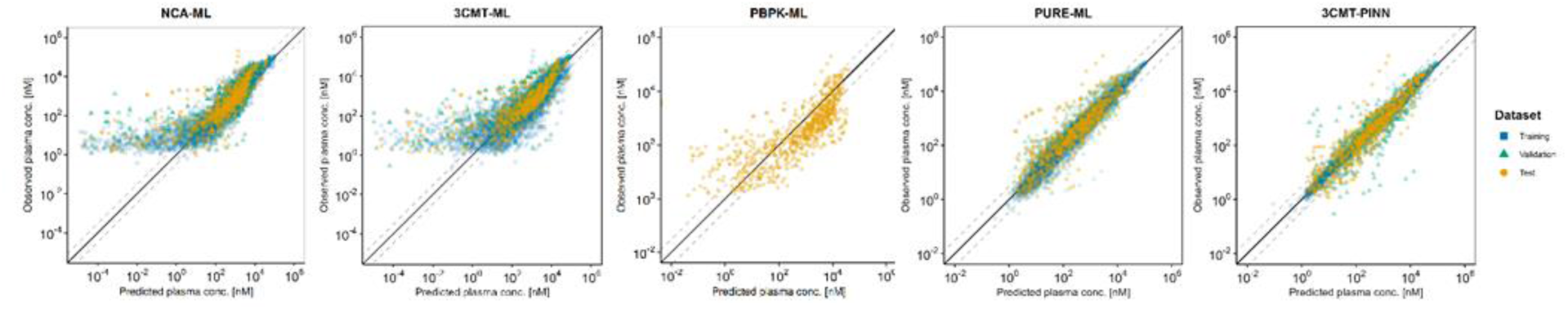
Predicted vs observed concentration data points separated into training, validation and test data sets across the different approaches.

**Table 3:** Percentage of predicted concentration data points within a two-fold and three-fold error compared to observed mean concentration data points for the test data set across the different approaches.

| Approach | NCA-ML | PBPK-ML | 3CMT-ML | PURE-ML | 3CMT-PINN |
| --- | --- | --- | --- | --- | --- |
| **2-fold** | 11.6% | 18.6%-27.8% | 8.98% | 61.0% | 65.9% |
| **3-fold** | 18.0% | 30.9%-45.1% | 14.3% | 79.7% | 83.5% |

For the specific PK metrics, the median relative errors for AUC, C_0_, and C_min_ (Table 4) revealed that NCA-ML, PBPK-ML, and 3CMT-ML performed poorest. NCA-ML and 3CMT-ML generally underpredicted these parameters with high relative errors (NCA-ML: 1.45/3.88/5.67, 3CMT-ML: 2.46/1.76/44.6), while PBPK-ML tended to overpredict AUC and C_0_ with values ranging from 0.31-0.40 for AUC, 0.29-0.97 for C_0_, and 0.05-1.97 for C_min_. In contrast, PURE-ML (1.20/1.36/0.796) and 3CMT-PINN (1.14/1.12/0.635) showed comparable results with values closer to the ideal value of 1, with the greatest difference seen in C_0_ predictions.

**Table 4:** Median of relative errors between observations and predictions for AUC, C0 and Cmin for the test data set across the different approaches.

| Approach | AUC<br>Median<br>Rel. error | C0<br>Median<br>Rel. error | Cmin<br>Median<br>Rel. error |
| --- | --- | --- | --- |
| **NCA-ML** | 1.45 | 3.88 | 5.67 |
| **PBPK-ML** | 0.31-0.40 | 0.29-0.97 | 0.05-1.97 |
| **3CMT-ML** | 2.46 | 1.76 | 44.6 |
| **PURE-ML** | 1.20 | 1.36 | 0.796 |
| **3CMT-PINN** | 1.14 | 1.12 | 0.635 |

**Table 5:** Median and percentage of geometric mean fold error (GMFE) lower than 2,3,5,10 and 100 between estimated and predicted concentration data across the different approaches.

| Test-Set | NCA-ML | PBPK-ML | 2CMT-ML | 3CMT-ML | PURE-ML | 3CMT-PINN |
| --- | --- | --- | --- | --- | --- | --- |
| **Median GMFE** | 8.15 | 3.51-7.80 | 20.8 | 7.36 | 2.09 | 2.10 |
| **% GMFE < 2** | 15.7 | 8.57-21.4 | 12.1 | 18.8 | 45.9 | 46.4 |
| **% GMFE < 3** | 24.3 | 18.6-40.0 | 19.7 | 29.0 | 65.2 | 72.5 |
| **% GMFE < 5** | 41.4 | 40.0-58.6 | 33.3 | 34.8 | 81.2 | 88.4 |
| **% GMFE < 10** | 55.7 | 55.7-80.0 | 45.5 | 53.6 | 85.3 | 97.1 |
| **% GMFE < 100** | 78.6 | 81.2-98.6 | 57.6 | 71.0 | 95.3 | 100 |

In addition to the derived error margins on which the ML methods were trained and derived PK metrics, a visual inspection of the resulting shape of the predicted PK profiles is used as additional quality measure. The visual representation of predicted PK profiles illustrates how well the overall profile shape and thereof derived PK metrics AUC, C_0_, and C_min_ were predicted. The closer the predicted profile approximates experimental data, in particular the observed C_max_ and C_min_ values and the segue between them, the better thereof derived PK metrics are predicted. Figure 3 shows representative PK profiles good (error or metric), moderate (error or metric), and bad (error or metric) predictions with the first row focusing on early (up to 8 hours) and the second row on terminal sampling points (until 24 hours) after drug administration. The shaded area represents the convex hull of the PK profile predictions from the ten different PBPK-ML models illustrating the bounds of model-predicted exposure.

**Figure 3:**
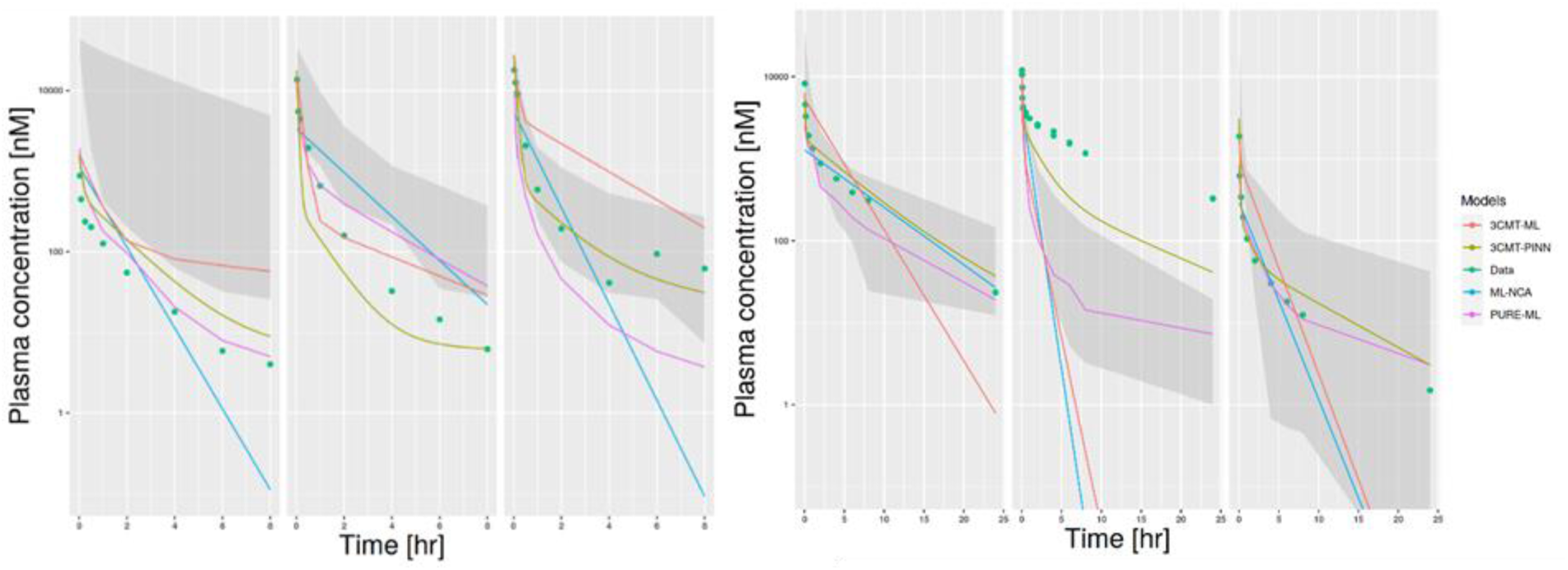
Predicted concentration-time profiles vs. observed mean concentration-time data for a good, moderate and bad prediction separated into PK profiles with tlast=8 and 24. Shaded area represents the convex hull of the PK prediction profiles from the 10 PBPK-ML models.

Comparing the training of NNs on plasma concentration-time data in the PINN setting to the training on derived PK parameters shows that the PINN approach performs better in predicting the concentration-time profiles. The CMT-PINN approach showed superior performance metrics and minimal GMFEs compared to alternative approaches, with the PURE-ML approach exhibiting comparable but showing slightly lower prediction accuracy across the different evaluation criteria. The substantial performance gap between these two approaches and the other methods (NCA-ML, CMT-ML, and PBPK-ML) highlights the advantage of PINNs for PK profile prediction.

## DISCUSSION

In this study, we conducted an objective comparison of five distinct ML methods (NCA-ML, PBPK-ML, CMT-ML, PURE-ML, and CMT-PINN) for predicting *in vivo* plasma concentration-time profiles from molecular structures. By using the same underlying *in vivo* PK dataset and split for all approaches, we enabled systematic benchmarking under identical conditions. The CMT-PINN approach achieved the highest prediction accuracy across evaluation metrics, with PURE-ML producing comparable but generally less accurate predictions. Our analysis yielded two principal findings: (1) The CMT-PINN methodology demonstrated superior performance for predicting PK profiles, exhibiting the highest accuracy across the examined evaluation metrics (2) Model training directly on concentration-time profiles rather than utilizing pre-calculated parameters resulted in enhanced predictive accuracy, with particularly pronounced improvements observed for the chosen small sample size scenario.

The literature on PK prediction methodologies has expanded considerably in recent years, demonstrating the feasibility of predicting PK metrics and profiles based solely on chemical structures (Table 1). These publications encompass both single-method innovations and systematic comparisons of multiple methodologies aimed at identifying optimal predictive performance.

Becker et al. recently demonstrated good prediction accuracy using PINNs, though without comparing to other model-based ML methods.^19^ The presented work extends their findings by benchmarking PINNs against both pure and hybrid ML methods using identical underlying data, confirming that PINNs trained directly on PK plasma profiles show advantages over approaches trained on derived NCA or estimated compartmental model parameters.

Contrary to our expectations, the use of two or three compartmental models with ML-predicted PK parameters (CMT-ML) did not yield better results compared to the reference (NCA-ML) approach, despite having more degrees of freedom to predict multiphasic PK profiles. The poor prediction accuracy of the PK parameters (NCA-ML mean-Pearson r² score: 0.68 vs. 2CMT-ML: 0.54, 3CMT-ML: 0.38) propagated to lower prediction accuracy of the concentration-time profiles. These results indicate the limitation of PK parameter based profile prediction methods on sparse datasets which was not observed in related studies with larger data sets.^27^

Walter et al. proposed the GMFE as a metric that does not bias toward early PK sampling time points.^27^ Using this metric, our predictions based on NCA yielded similar GMFE values to those reported by Walter et al., with improved results when training was performed directly on concentration-time profiles rather than NCA parameters. The compartmental model approach (CMT-ML) performed better in Walter et al. compared to our analysis, indicating again that PK parameter based methods require larger data sets and compound diversity compared to the superior performing CMT-PINN approach. Based on GMFE values, the CMT-PINN approach performed best across all tested methods with 5-21 percentage points higher GMFE values. Similarly, when comparing the percentage of predicted concentration-time values within 2-fold and 3-fold ranges.

Compared to Beckers et al.,^19^ we demonstrated that the CMT-PINN approach remains effective even when selecting a smaller dataset for training (selected ∼700 studies versus 13,000 reported experiments). Our neural network architecture is more compact, employing only 3 instead of 5 fully connected layers, half the hidden units, and no additional penalty for higher compartments. These adaptations might be better suited for our selected smaller data set. Here a systematic analysis relating increasing number of studies for training and varying the selected chemical space could help to explore the relationship between model architecture, prediction accuracy, and selected data sets.

Multiple studies evaluated PBPK-ML approaches with varying degrees of success. ^10,18,27^ While the PBPK-ML approach gives comparable results to other approaches, its performance heavily depends on the quality of underlaying properties and parameter measurements, accuracy of derived *in vitro* parameter predictions, the level of detail of the used active processes, and their contextualization within the PBPK framework. Notably, even the additional optimization of critical model parameters lipophilicity and clearance did not result in outperforming prediction accuracy of the reported PBPK-ML approach.^27^ This underlines the need for a systematic incorporation of detailed ADME processes within the PBPK framework rather than building on physicochemical properties and unspecific liver processes as driving properties for plasma PK predictions. However, data required for predicting the mechanisms of ADME processes early in drug discovery is still scares and most PBPK applications occur during development stages where most of the ADME processes are already well characterized.^42^

Another promising method, which has not yet been tested in the context of predicting PK profiles based on molecular structure are universal differential equations (UDEs) and neural ordinary equations. A recent example used UDEs to study the absorption rate constant and nonlinear clearance in a human case example.^43^

## OUTLOOK

The systematic evaluation of predictive methodologies presented in this work establishes a framework for rigorous assessment of emerging approaches in PK prediction. Rather than conducting isolated proof-of-concept studies with limited industrial impact, our approach enables comprehensive comparison of prediction methodologies on consistent datasets, providing unbiased assessment of their predictive performance.

To facilitate adoption of these computational methods in drug discovery workflows, several key developments might be useful.

1. Enhanced uncertainty quantification: future implementations should incorporate robust uncertainty quantification for predictions, allowing researchers to make informed decisions based not only on predicted values but also on confidence intervals.^44,45^
2. Defined applicability domains: establishing clear applicability domains for these models is essential to ensure reliable predictions for novel chemical structures and to prevent misleading extrapolations beyond the model’s reliable prediction space.^45,46^
3. User-friendly implementation: successful integration into day-to-day decision-making processes requires user-friendly applications that provide transparent and educative supplementary information beyond mere predictive outputs.
4. Expanded mechanistic understanding: future models should better incorporate mechanistic understanding of non-linear ADME processes to improve predictions for compounds with complex PK behaviors such as time-dependent auto-induction or inhibition of metabolizing enzymes.
5. FAIR AI datasets and models: implementation of Findable, Accessible, Interoperable, and Reusable (FAIR) principles for both datasets and models is crucial for advancing the field. Expanding from simple result metrics to comprehensive and standardized metadata, clear documentation of data provenance, interoperable data formats, and version-controlled model repositories are required.

Implementing the best-performing prediction methodologies as new services for research teams and computational scientists constitute a pivotal advancement toward the “in silico first” paradigm based on data-driven decision making. This approach will continuously innovate design-make-test-analyze (DMTA) cycles. Over time, such validated and systematically applied computational predictions will increasingly supersede conventional experimental procedures, accelerating compound synthesis selection, reducing *in vitro* assay screenings and *in vivo* animal studies, and shortening overall research timelines.

This systematic approach of developing, evaluating, and implementing the best preforming prediction methods in continuous DMTA cycles builds towards an “Applied AI Factory” with accelerated decision making for compound synthesis, research timelines, reduce animal studies and ultimately bringing the right medicine faster and as a commodity to patients in need.

## ACKNOWLEDGEMENTS

We would like to thank our Sanofi colleagues who designed and synthesized the compounds, executed the in vitro & in vivo experiments, conducted the sample measurements & analysis, and transferred their results into Sanofi’s F.A.I.R. data environments that enabled this work. Further we would like to acknowledge Julius Martensen for his helpful discussions on PINNs and their implementation in Julia. Also, we would like to acknowledge Jannis Schuhmann for his contributions to the hyperparameter optimization of the XGBoost model during his internship.

## AUTHOR CONTRIBUTIONS

F.J. and H.C. conceptualized and designed the research; F.J., C.G., H.M. and H.C performed the research and analyzed the data; F.J., C.G., H.M. and H.C wrote the manuscript.

## SUPPLEMENT

**Table S1:** Number of PK parameter sets separated into one, two and three compartmental models depending on coefficient of variation (CV) filtering. Filtering ensures that all estimated parameters have CVs lower than a certain percentage.

| **CV [%]** | 10 | 20 | 30 | 40 | 50 | 60 | **70** | 80 | 90 | 100 |
| --- | --- | --- | --- | --- | --- | --- | --- | --- | --- | --- |
| **1CMT** | 77 | 130 | 173 | 187 | 187 | 190 | **190** | 176 | 172 | 166 |
| **2CMT** | 33 | 149 | 197 | 252 | 295 | 317 | **335** | 348 | 356 | 357 |
| **3CMT** | 4 | 18 | 44 | 81 | 105 | 123 | **132** | 148 | 159 | 171 |
| **# models** | 114 | 297 | 414 | 520 | 587 | 630 | **657** | 672 | 687 | 694 |

